# Endogenous and exogenous control of visuospatial selective attention in freely behaving mice

**DOI:** 10.1101/550822

**Authors:** Wen-Kai You, Shreesh P. Mysore

## Abstract

Selective spatial attention, the ability to dynamically prioritize the most important spatial location, is essential for adaptive behavior. It has been studied primarily in head-fixed animals, and almost exclusively in primates. Here, we report the development of two human-inspired, discrimination-based behavioral paradigms for studying selective visuospatial attention in the freely behaving mouse: the spatial probability task, and the flanker task. In the spatial probability task, we found enhanced response accuracy, perceptual discriminability, and rates of sensory evidence accumulation at the location with higher probability of target occurrence, and opposite effects at the lower probability location. In the absence of systematic differences in sensory input, motor biases, and trial structure, these results demonstrated endogenous expectation-driven shifts of spatial attention. In the flanker task, we found that a second, ‘flanker’ stimulus presented with the target, but with incongruent information, caused switch-like decrements in response accuracy and perceptual discriminability as a function of flanker contrast, as well as a reduced rate of evidence accumulation. These results demonstrated exogenous capture of spatial attention. The innovation of behavioral tasks for selective visuospatial attention in unrestrained mice opens up a rich avenue for future research dissecting the neural circuit mechanisms underlying this critical executive function.

## INTRODUCTION

Animals have the remarkable ability to preferentially process the most important, or “highest priority”, information in complex environments to guide behavior. Called selective attention, this ability is essential for a range of cognitive functions and adaptive behavior, and its dysfunction is found in diverse psychiatric illnesses including ADHD and schizophrenia. A rich body of work into visual selective attention has provided insights into the consequences of selective attention to behavior and neural processing (Carrasco, 2011), and has identified the involvement of critical fronto-parietal (Bisley and Goldberg, 2010; Squire et al., 2013) and midbrain (Krauzlis et al., 2013) networks in selective attention. However, the neural circuit mechanisms for the control of selective visuospatial attention remain largely open questions.

The dissection of these circuit mechanisms, including those underlying space-specific target selection and distracter suppression, can benefit greatly from cutting-edge approaches that allow for the interrogation of subsets of neurons that are distinguished by their shared functional identity, shared patterns of anatomical connectivity, and so on. Such approaches necessitate the use of a genetically tractable animal system. In contrast to this need, visual selective attention has, thus far, been studied nearly exclusively in primates (but see (Sridharan et al., 2014) and (Wang and Krauzlis, 2018)), in which such genetic approaches are difficult to implement. Whereas the mouse, a genetically powerful animal model, has been used to study visually guided behavior (Burgess et al., 2017; Busse et al., 2011; Bussey et al., 2001; Itokazu et al., 2018; Marques et al., 2018), and recently, selection between visual and other sensory stimuli (Wimmer et al., 2015), their lower visual acuity compared to primates (Histed et al., 2012; Prusky and Douglas, 2004), and their inherent impulsivity (Isles et al., 2004), have led to the concern that mice may not be an effective choice for studying higher order visual cognitive function (Huberman and Niell, 2011). Specifically, it is debated whether mice are capable of exhibiting spatially well-resolved visual behaviors that are necessary to unpack the circuit basis of visuospatial attention, with just one recent study reporting success (Wang and Krauzlis, 2018).

Second, because selective attention operates in nature in animals engaging freely with the world, the study of its neural underpinnings can benefit greatly from investigations in unrestrained animals in a behavioral state resembling natural conditions: with intact vestibular feedback cues (Rancz et al., 2015) as well as coordinated body movements (Aghajan et al., 2015; Chen et al., 2013; Ravassard et al., 2013). Many reports have highlighted differences in neural representations as a function of the behavioral state of the animal (Cardin and Schmidt, 2004; Cohen and Castro-Alamancos, 2010; Fu et al., 2014; Keller et al., 2012), and there has been a recent push towards measuring behavioral parameters including head and eye positions in unrestrained animals (Meyer et al., 2018; Payne and Raymond, 2017). In contrast to these recent directions, selective (visuospatial) attention has been studied exclusively in head-fixed preparations.

Therefore, the ability to study selective visuospatial attention in freely behaving animals in which diverse genetic tools may be brought to bear would offer a powerful advantage for the deconstruction of the neural basis of naturalistic visuospatial attention control. More generally, the need for well-characterized behavioral paradigms has recently been highlighted as a critical building block for circuit neuroscience (Krakauer et al., 2017). In response to these needs, we present, for the first time, human-inspired behavioral tasks for the study of endogenous as well as exogenous control of visuospatial selective attention in freely behaving mice. Both tasks described in this study are touchscreen-based, self-paced, and dissociate the locus of spatial attention from the locus of behavioral report.

## RESULTS

All the behavioral tasks in this study involved a touchscreen-based (Bussey et al., 2001; Morton et al., 2006) setup (Methods). Freely behaving mice were placed in a plexiglass tube within a soundproof operant chamber equipped with a touch-sensitive screen, and a reward well located at the opposite face of the box from the touchscreen (Fig. 1A). A plexiglass sheet, with three holes corresponding to the locations at which the mouse was allowed to interact with the touchscreen by a nose-touch, was placed in front of it. All trials began with a nose-touch on a bright zeroing-cross presented within the lower central hole, following which visual stimuli (bright objects on a dark background) were presented elsewhere on the screen. The upper holes served as response ports for the animals to report their behavioral choice. Behavioral data was collected from daily sessions that lasted 30 minutes for each mouse.

**Figure 1.**
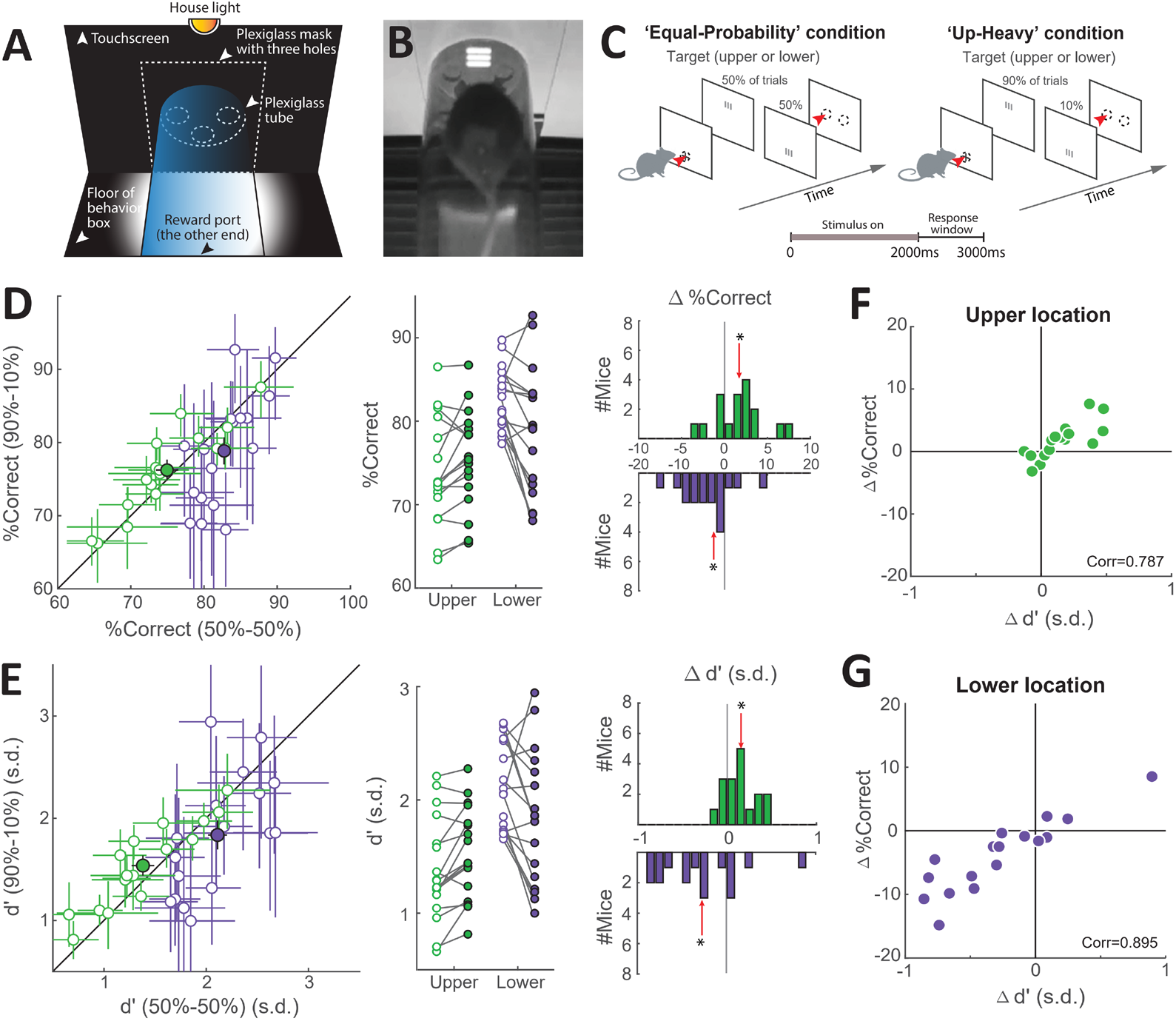
Spatial probability modulates response accuracy and perceptual sensitivity in a space-specific manner in freely behaving mice. **(A)** Schematic touchscreen-based setup for visually-guided tasks with unrestrained mice, showing key components. **(B)** Snapshot of mouse facing a visual stimulus on the touchscreen. **(C)** Schematic of task design. Trials began with a nose-touch (red arrowhead) on a zeroing-cross presented within the lower central hole (dashed oval). A single oriented grating (’target’) was presented after trial initiation (size= 25o, 60 pixels x 60 pixels; duration = 2s, contrast=15.2%; Methods). <u>Left panel</u>: Equal probability (’50-50’ condition). Target waspresented with equal probability at the upper or lower locations (Methods). <u>Right panel</u>: Up-heavy (’90u-10’condition). Target was presented with 90% probability at the upper location. Mice were rewarded for reporting the orientation of the target grating: vertical → nose touch to the left; horizontal → nose touch to the right. The two conditions were run in blocks (one per day), pseudo-randomized across days. Dashed ovals: response holes; the ‘zeroing’ hole is shown only in the first screen, and the two response holes are shown only in the third screen for clarity. Red arrowheads denote nose-touches. **(D)** Effect of spatial probability on response accuracy (n=17 mice; Fig. S1D; Methods). <u>Left panel:</u> Scatter plot comparing the performance of mice in the 90u-10 conditionagainst the 50-50 condition; intended for visualization of variability in the dataset. Green: upper location, purple: lower location; error bars: 95% confidence intervals (obtained by bootstrapping; Methods); filled dots: group mean values ± S.E.M. <u>Middle panel</u>: Data replotted to compare mean response accuracy (% correct) ofeach mouse (line) between the 50-50 (hollow dots) and 90u-10 (filled dots) conditions. <u>Right panel</u>: Distributionsof change in accuracy (90u-10 condition minus 50-50 condition) at the upper location (top row) and lower location (bottom row); derived from middle panel. Red arrows: median; * p<0.05; signed-rank and HB tests (Methods). **(E)** Effect of spatial probability on perceptual sensitivity (d’); conventions as in D. **(F)(G)** Change in accuracy plotted against change in d’ at the upper (F) and lower (G) locations; data show positive correlation; corr = Pearson’s correlation coefficient. See also Figure S1 and Video S1.

### Mice learn feature-response association rule through a single-stimulus visual discrimination task

A key aspect of the design of our tasks for spatial attention involved the decoupling of the spatial locus of the target of attention from that of the behavioral report. To achieve this, all mice were first trained to generate behavioral responses based on feature information contained in the target stimulus – here, orientation of the target grating – rather than based on its spatial location (Fig. S1A). Immediately upon trial initiation (nose-touch at the zeroing cross), a single oriented grating (‘target’) was presented at a fixed location along the vertical midline (Methods). Mice were rewarded if they responded to a vertically oriented target with a nose-touch to the left response port, and to a horizontally oriented grating with a nose-touch to the right response post (Fig. S1A).

Mice can discriminate orientations of visual gratings (Andermann et al., 2010; Long et al., 2015; Wang and Krauzlis, 2018), and, consequently, learned well the response association rule (Fig. S1B-D; Methods). The highest accuracy across mice trained on this task was 94.5%, and the median accuracy was 84.3% with a 95% confidence interval of [81.6%, 87.1%] (n=20 mice; Fig. S1BD). The median values of the response reaction times (RTs) of these mice were distributed between 480 ms and 920 ms, with a median value of 700 ms (Fig. S1CD).

To examine if there were any systematic biases in mouse performance, we tested for differences in the responding of mice to horizontal versus vertical gratings. Response accuracy and RTs to horizontal gratings were not different from those to vertical gratings (Fig. S1EF; Wilcoxon signed-rank test followed by the Holm-Bonferroni test for multiple comparisons, referred to as ‘signed-rank and HB tests’ henceforth; Methods), nor were the RTs to left versus right nose-touches (Fig. S1G; signed-rank test). These results showed that there were no systematic sensory or perceptual biases with respect to grating orientation, nor motor biases with respect to nose-touch location.

Thus, mice were able to learn a fixed association rule between target orientation and a distinct response location, thereby dissociating successfully the loci of sensory input and motor output. With this task as a foundation, we next trained freely behaving mice on selective spatial attention tasks.

### Endogenous (top-down) control of selective visuospatial attention

To study endogenous control of visuospatial attention, we trained freely behaving mice on a spatial probability task (Fig. 1C). Previous work in humans has shown that manipulating the spatial probability of target occurrence can serve as an endogenous attentional cue (Druker and Anderson, 2010; Geng and Behrmann, 2005; Vincent, 2011). Here, immediately upon trial initiation, a single oriented grating (‘target’) was presented on the screen at one of two locations along the vertical axis. One location was far above the zeroing cross and referred to as the ‘upper’ location, while the other was just above the zeroing cross and referred to as the ‘lower’ location (Fig. 1BC; Methods). Mice were rewarded for correctly reporting the orientation of the target (presented at one of two locations along the vertical axis) with an appropriate nose-touch (at one of two locations along the horizontal axis), per the association rule learned previously.

The probability of target occurrence at the upper and lower locations was changed in different blocks of trials in order to manipulate the mouse’s expectation regarding target location. The blocks were of two types: (i) Equal probability (or ‘50-50’) block, in which, on each trial, the target occurred with equal (50%) probability at the upper or the lower locations (Fig. 1C; left panel), and (ii) Up-heavy (or ‘90u-10’) block, in which, on each trial, the target occurred with 90% probability at the upper location, and 10% probability at the lower location (Fig. 1C; right panel). Each block lasted for the entire behavioral session on a given day (30 minutes), and blocks of the two types were interleaved randomly across days.

In the 90-10 block, the upper location was chosen to be the higher probability location because pilot experiments revealed an asymmetry in perceptual performance between the two stimulus locations. In the baseline 50-50 condition, response accuracy at the upper location was consistently worse than at the lower location (Fig. S1H, median difference =-10.9%, p<0.001, signed-rank test). Therefore, biasing the probability of target occurrence in favor of the upper location allowed us to test if biased spatial expectation could improve behavioral performance, without any potential confounds due to ceiling effects.

To examine the effects of spatial probability of the target on behavioral performance, we started by comparing the response accuracy of mice in the 90u-10 versus the 50-50 blocks for each target location. To this end, for each mouse, we pooled all trials in which the target was at the upper location, and separately all trials in which the target was at the lower location. For each location, we then compared the %-correct values between block types (Figs. 1D, S1I; Methods).

We found that at the upper location, mice (n=17) exhibited a significant improvement in response accuracy in the 90u-10 blocks over that in 50-50 blocks (Fig. 1D; green data, median improvement = 1.9%; p=0.028, signed-rank and HB tests; Methods). By contrast, at the lower location, we found that animals exhibited a significant worsening of accuracy in the 90u-10 blocks (Fig. 1D, purple data; median reduction = 2.5% improvement; p=0.013, signed-rank and HB test). (Thus, taken together, the spatial probability manipulation yielded a net swing in performance of 4.4% between the upper and lower locations; Fig. 1D; right column.) Thus, changes to spatial probability of the target modulated performance in a spatially selective manner favoring the higher probability location.

Next, we examined the effect of spatial probability on two independent factors that impact response accuracy, namely, perceptual sensitivity (d’) and decision criterion (c) (Stanislaw and Todorov, 1999). Heightened perceptual sensitivity (d’) improves accuracy, as does reduction in the magnitude of criterion (c), whereas reduced d’ or an increase in |c| worsens accuracy (Fig. S1J). Using ideal observer analysis, we computed d’ and c at each of the two locations, and for each block-type.

We found that at the upper location, animals exhibited an increase in perceptual sensitivity in 90u-10 blocks compared to 50-50 blocks (Fig. 1E; green data; median increase=+0.14 s.d., p=0.006, signed-rank and HB tests; Methods). By contrast, at the lower location, animals exhibited a significant decrease in perceptual sensitivity (Fig. 1E, purple data; mean decrease=-0.3 s.d., p=0.028, signed-rank and HB tests). (Taken together, the spatial probability manipulation yielded a net swing in d’ of 0.44 s.d. between the upper and lower locations; Fig. 1E; right panel.) There was no systematic effect of spatial probability on the decision criterion at either location (Fig. S1K; upper location (green data): p=0.981; lower location (purple data): p=0.332, signed-rank and HB tests). Indeed, at both locations, changes in accuracy correlated well with changes in perceptual sensitivity (Fig. 1F, upper location, Pearson correlation coefficient=0.787, p<0.001; Fig. 1G, lower location, Pearson correlation coefficient=0.895, p<0.001), but not with changes in criterion (Fig. S1L, upper location, green data: Pearson correlation coefficient=-0.271, p=0.292; lower location, purple data: Pearson correlation coefficient=-0.140, p=0.592), indicating that changes in sensitivity, but not criterion, best accounted for the effects on accuracy. Thus, changes to spatial probability of the target also modulated perceptual sensitivity of mice in a spatially selective manner, favoring the higher probability location.

We next investigated the effect of spatial probability on the RTs of mice in this task. As a first step, we compared the median RTs of mice in the 90u-10 block versus the 50-50 block at each target location. We found that at the upper location, animals exhibited faster RTs during 90u-10 blocks versus 50-50 blocks (Fig. 2A, green data; median change=-20 ms, p=0.031, signed-rank and HB tests), consistent with a potential higher expectation of target occurrence. Surprisingly, we found faster RTs at the lower location as well (Fig. 2A, purple data; median change=-50 ms, p=0.002, signed-rank and HB tests).

**Figure 2.**
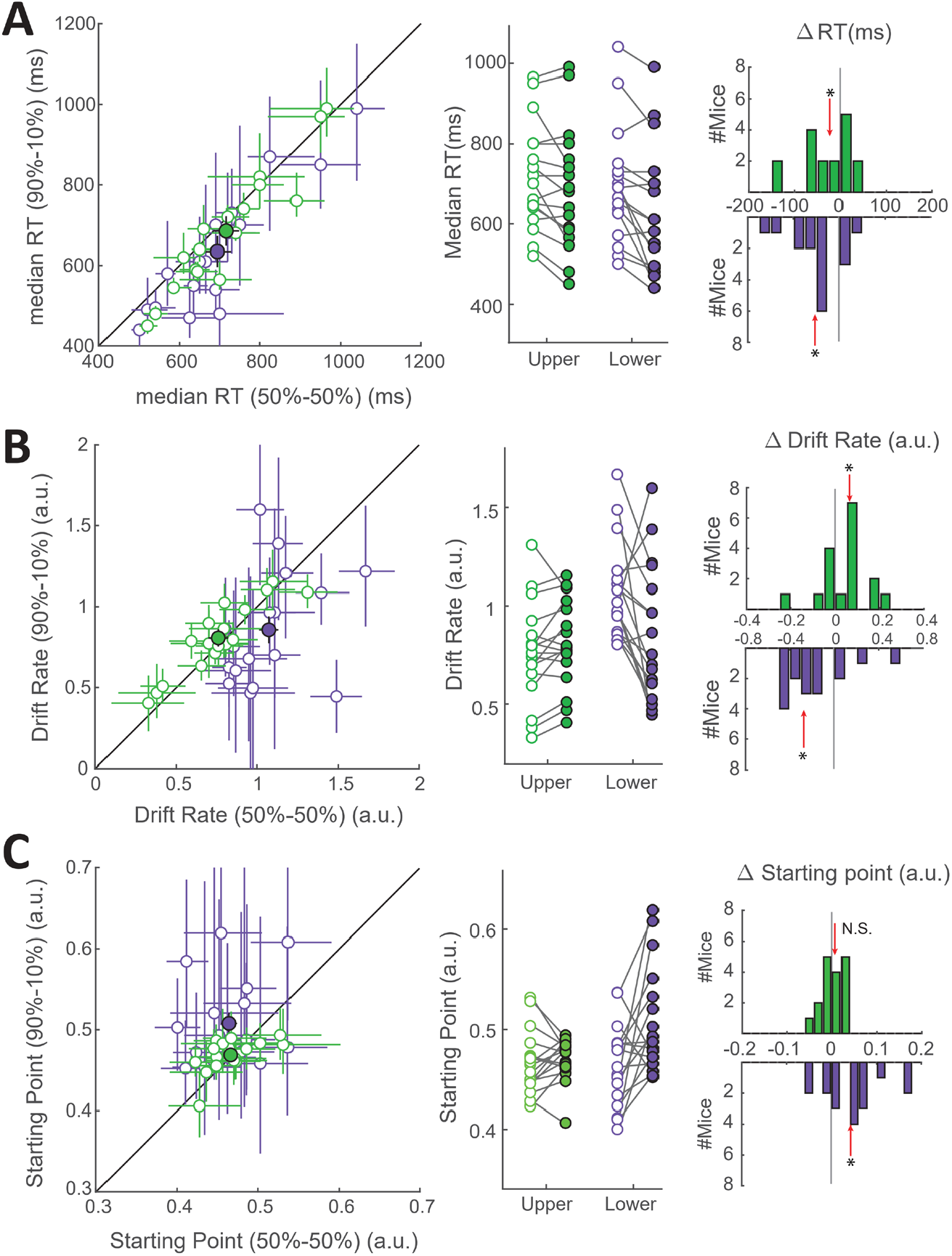
Spatial probability modulates rate of sensory evidence accumulation in a space-specific manner in freely behaving mice. **(A)** Effect of spatial probability on median reaction time (RT) (n=17 mice); conventions as in Fig. 1DE. <u>Leftpanel:</u> Scatter plot comparing the median RT of mice in the 90u-10 condition against the 50-50 condition.Green: upper location, purple: lower location; error bars: 95% confidence intervals (obtained by bootstrapping; Methods); filled dots: group mean values ± S.E.M.<u>Middle panel</u>: Comparison of median RT of each mouse (line) between the 50-50 (hollow dots) and 90u-10 (filled dots) conditions. <u>Right panel</u>:Distributions of change in median RT (90u-10 condition minus 50-50 condition) at the upper location (top row)and lower location (bottom row). Red arrows: median; * p<0.05; signed-rank and HB tests (Methods). **(B-C)** Drift diffusion modeling of RT distributions from spatial probability task; data from each mouse were fitted to the 4-parameter model, and the best fit values of the parameters were obtained for each mouse (Methods; Fig. S2). Shown are effects on two of the four parameters: rates of sensory evidence accumulation (drift rates, B), and starting point (C). Conventions as in A. See also Figure S2.

To gain insight into these effects, we turned to diffusion modeling of RTs, which considers the full distribution of observed RTs and unpacks potential underlying factors that explain them (Ratcliff, 1978; Voss et al., 2013). Since we observed no systematic biases in perceptual performance w.r.to vertical versus horizontal targets (Fig. S1E-G), we pooled both target types together, and fit the RT distributions of correct vs. incorrect trials with a two-choice drift-diffusion model (Methods). This approach allowed us to estimate for each mouse, and at each target location, the rate of evidence accumulation (drift rate), the separation between decision boundaries for correct and incorrect responses, the starting point of evidence accumulation, and the non-decisional constant (t0) in each block type (Fig. S2A). To obtain accurate estimates of these model parameters, trials with outlier values of RTs are typically dropped before drift-diffusion modeling (Voss et al., 2015), and we used a bootstrapping-based approach to identify trials with inordinately short or long RTs for exclusion (Fig. S2B; Methods). For the following analysis, only trials with RTs between 300ms and 2250 ms were included, representing 96% of total trials; outlier trial exclusion did not alter the effects on median RT in Figure 2A (Fig. S2C).

We found that when the target was at the upper location, the drift rate was faster in 90u-10 blocks relative to 50-50 blocks (Fig. 2B, green data, median change=+0.06 a.u., p=0.035, signed-rank and HB tests), indicating a faster rate of evidence accumulation. None of the other parameters were significantly different at the upper location between the two block types (Fig. 2C, S2DE; green data). By contrast, when the target was at the lower location, the drift rate was slower in 90u-10 blocks relative to 50-50 blocks (Fig. 2B, purple data, median change=-0.26 a.u., p=0.017, signed-rank and HB tests), indicating a slower rate of evidence accumulation. In addition, the starting point was higher, i.e., biased towards the correct response (Fig. 2C, purple data, median change=+0.04 a.u., p=0.017, signed-rank and HB tests), and the separation between decision boundaries exhibited a trend towards smaller values (Fig. S2D, purple data, median change=-0.06 a.u., p=0.039, not significant per signed-rank and HB tests). These observations indicated that, in the 90u-10 blocks, the threshold of evidence needed to trigger correct responses when the target was at the lower location was significantly reduced (Fig. S2F; purple data).

These results revealed two key insights. Firstly, that in the 90u-10 blocks, the rates of evidence accumulation at the two target locations were affected in opposite ways: faster drift rates at the upper location, consistent with a potential increase in spatial expectation of target, and slower drift rates at the lower location, consistent with a potential decrease in spatial expectation. Secondly, that the speeding up of RTs observed at both the upper and lower locations in the 90u-10 blocks were driven by different factors. At the upper location, faster reaction times in the 90u-10 blocks were consistent with the faster drift rates. By contrast, at the lower location, faster reaction times despite slower drift rates were consistent with the reduced thresholds for triggering correct responses. If true, this lower threshold, i.e., the less stringent requirement for sensory evidence (or ‘overconfidence’), would predict more errors when the target was at the lower location, a prediction that matched our observations (Fig. 1D, purple data).

Taken together, our results demonstrated space-specific effects of spatial probability on behavioral performance: compared with the 50-50 blocks, the 90u-10 blocks showed (1) increased accuracy, (2) improved perceptual sensitivity, and (3) faster rate of evidence accumulation at the upper location –i.e., the target location with higher spatial probability; and a concurrent decrease in all three metrics at the lower location.

These observed effects occurred in the absence of any systematic differences in sensory input between block-types at the two target locations (Fig. S1M). They occurred also in the absence of any changes in motor biases toward particular response locations (Fig. S1N). There were also no systematic differences in local trial structure because of the design of the task: when the target appeared at the same location in successive trials, say the upper location, there was no difference between the probabilities that the two targets were the same versus that they were different (both 50%), nor any difference in these probabilities between the 90u-10 and 50-50 conditions. Additionally, when the same target appeared at a particular location in two successive trials, say the upper location, the probability of a correct response on the second trial was no different between the 90u-10 and 50-50 conditions (Fig. S1Q). Together, these results ruled out sequential effects or visual after-effects (Quinlan and Hill, 1999). Finally, the long inter-trial interval (median = 16.4 s with 95% CI of [14.9 s, 17.9 s] in the 90u-10 condition, and median = 15.3 s with 95% CI of [13.8 s, 16.9 s] in the 50-50 condition), ruled out low level effects of repetition priming (Desimone, 1996; Hillstrom, 2000; Kristjansson et al., 2002; Maljkovic and Nakayama, 1996).

Consequently, the space-specific effects of spatial probability observed in the 90u-10 blocks were best explained by an expectation-driven shift of attention towards the upper location and away from the lower location, i.e., endogenously driven selective spatial attention.

### Exogenous (bottom-up) capture of selective visuospatial attention

To study exogenous control of visuospatial attention in freely behaving mice, we trained a majority of the same animals on a touchscreen version of the attentional flanker task used in humans (Eriksen and Eriksen, 1974; Fan et al., 2002) (Fig. 3AB; Methods). Here, immediately upon trial initiation, up to two oriented gratings were presented on the screen at two locations along the vertical axis. As with the spatial probability task, one location was far above the zeroing cross (‘upper’ location), and the other was just above the zeroing cross (‘lower’ location). The target grating, i.e., the one that yielded reward, was present on every trial, and *always* occurred at the lower location. The second, flanker grating, when present, *always* occurred at the upper location (Fig. 3AB). Mice were rewarded for reporting correctly the orientation of the target grating while ignoring that of the flanker (when also present).

**Figure 3.**
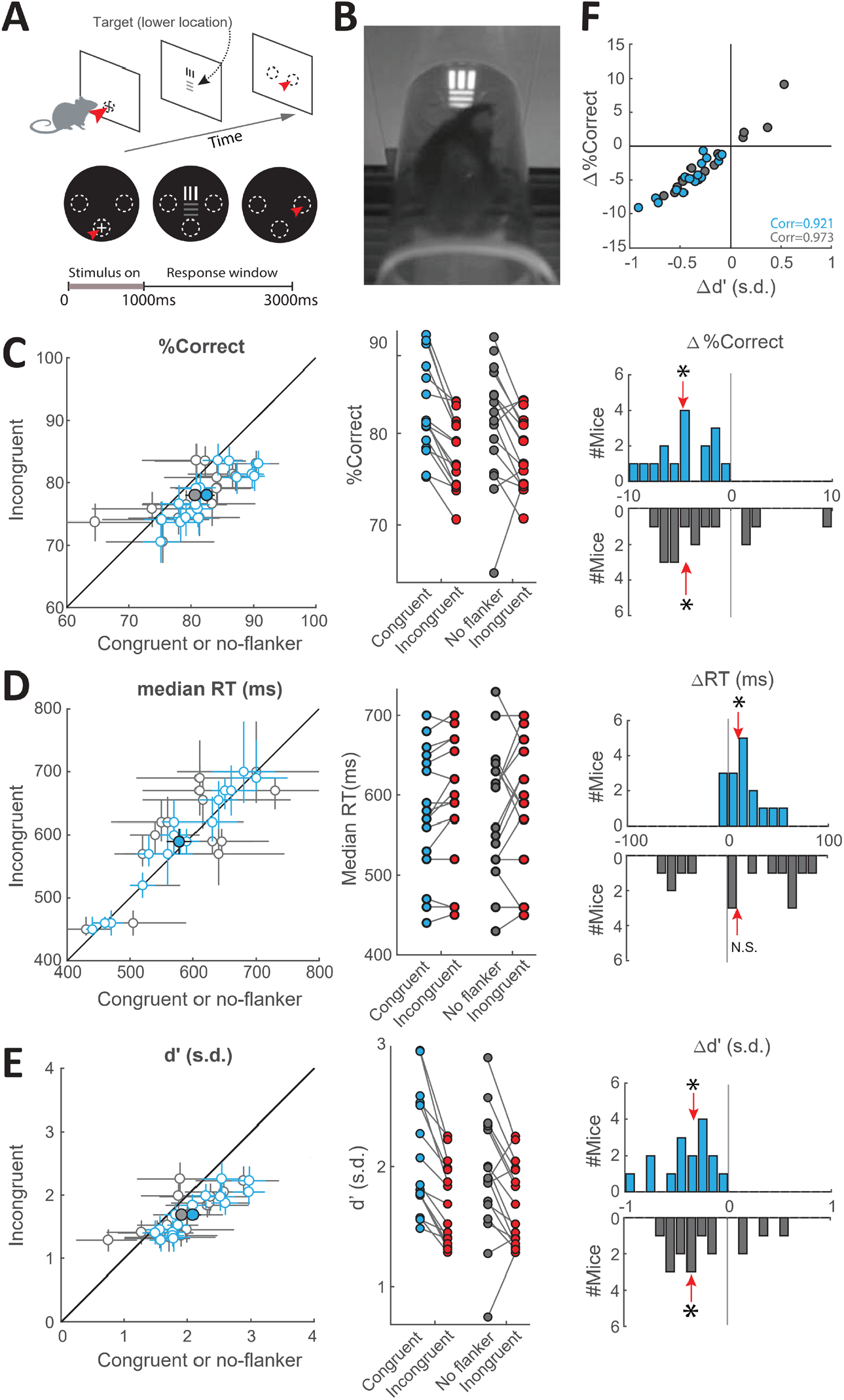
Incongruent flanker reduces response accuracy and perceptual discriminability, and increases RTs, in freely behaving mice. **(A)** Schematic of task design. <u>Top-row</u>: Following trial initiation, one or twograting stimuli were presented simultaneously. The target (one that yielded reward) was always presented at the lower location, and mice were trained to report the target’s orientation with an appropriate nose touch. A second stimulus (“flanker”), when present, was behaviorally-irrelevant, and was always presented at the upper location. Orientation of the flanker was either congruent (same) or incongruent (orthogonal) to the target across trials. Flanker contrast was varied parametrically in a randomly interleaved fashion across trials. <u>Bottom-row</u>: Screen-shots of display at different stages in an incongruent flanker trial. For this figure, data were analyzed by first collapsing across contrasts for each flanker condition. **(B)** Picture of mouse facing target and incongruent flanker. **(C)** Effect of incongruent flanker on response accuracy (n=16 mice); conventions as in Fig. 1.DE. <u>Left panel</u>: Scatter plot comparing response accuracy to target’s orientation in incongruent trials (red data) versus either the congruent trials (blue data), or versus the no-flanker trials (grey data). Large filled circles: group mean ± S.E.M. <u>Middle panel</u>: Comparison of mean response accuracy (%correct) of each mouse (line) between flanker conditions. <u>Right panel</u>: Distributions of changes in response accuracy; incongruent minus congruent (blue), or incongruent minus no-flanker (grey) trials. **(D-E)** Effect of incongruent flanker on median reaction time (D) and perceptual sensitivity (E); conventions as in C. **(F)** Plot of changes in response accuracy against changes in perceptual sensitivity. Blue data: incongruent trials versus congruent trials, grey data: incongruent trials versus no-flanker trials; corr=Pearson’s correlation; *: p<0.05. See also Figure S3 and Video S2.

Trials were of three types: (a) singleton trials, in which only the target grating was presented, (b) incongruent flanker trials, in which the flanker grating was also presented simultaneously, with the orientation of flanker being *orthogonal* to that of the target, and (c) congruent flanker trials, in which the orientation of the flanker was *identical* to that of the target. In both the incongruent and congruent flanker trial types, the physical salience of the flanker, here, its visual contrast, was varied across trials from being less than to being greater than that the contrast of the target. All three trial types, as well as the different contrasts of the flankers, were interleaved randomly in each behavioral session.

Studies in humans (Eriksen and Eriksen, 1974; Fan et al., 2002) show that incongruent flankers capture attention, effectively outcompeting the behaviorally relevant target frequently, and resulting in poorer performance (greater number of error trials). Informed by this, and the observed asymmetry in mouse perceptual behavior between the upper versus lower locations (Fig. S1H), we chose the lower location – the one with better perceptual performance– as the fixed location for the target stimulus, and the upper location as the fixed location for the flanker stimulus. This allowed us to test if a flanker affected behavioral performance in mice without a potential confound due to floor effects in performance. Additionally, our task design decoupled the location of the target stimulus from the locus of the behavioral report, and also permitted the parametric investigation of the effect of the flanker on performance.

As a first step in analyzing the results from mice trained on this task (n=16), we compared behavioral performance between incongruent and congruent flanker trials (Fig. 3C-E), and did so by collapsing data across flanker contrasts (Fig. S3A; Methods). We found that mice exhibited a significant impairment in response accuracy in the incongruent trials compared to the congruent trials (Fig. 3C, blue data; median change=-4.7%, p<0.001, signed-rank and HB tests), and a significant increase in the median RTs (Fig. 3D, blue data; median change=+10 ms, p=0.019, signed-rank and HB tests).

Upon partitioning response accuracy into perceptual sensitivity and decision criterion, we found a significant reduction in perceptual sensitivity in incongruent trials compared to the congruent trials (Fig. 3E, blue data; median change=-0.34 s.d., p<0.001, signed-rank and HB tests), and no change in decision criterion (Fig. S3B, blue data, p=1, signed-rank and HB tests). The observed reduction in response accuracy was correlated strongly with the reduction in perceptual sensitivity across animals (Pearson’s correlation coefficient = 0.921, p <0.001; Fig. 3F, blue data), and weakly with increases in the absolute value of criterion (Pearson’s correlation coefficient = −0.575, p =0.02; Fig. S3C, blue data).

Similar results were found upon comparing performance in the incongruent trials versus singleton (no-flanker) trials. We found a significant reduction in response accuracy (Fig. 3C, gray data; median change=-4.4%, p=0.026, signed-rank and HB tests) and in perceptual sensitivity (Fig. 3D, gray data; median change=- 0.35 s.d., p=0.023, signed-rank and HB tests) in the incongruent trials, with no change in decision criterion (Fig. S3B, gray data; p=0.438, signed-rank and HB tests). In addition, decreases in accuracy were strongly correlated with decreases in sensitivity across animals (Pearson’s correlation coefficient = 0.973, p<0.001, Fig. 3F, gray data), but not correlated with changes in criterion (Pearson’s correlation coefficient = 0.015, p=0.955, Fig. S3C, gray data). These results showed that in mice, as in humans (Eriksen and Eriksen, 1974; Fan et al., 2002), the incongruent flanker produced a reduction in accuracy and sensitivity, and a slowing down of reaction times, consistent with attention being captured by the flanker.

Next, we analyzed the dependence of these effects on the contrast of the incongruent flanker. We found that response accuracy in incongruent trials varied with flanker contrast in a striking manner (Fig. 4A, red data). As long as the contrast of the flanker was weak, response accuracy was not significantly different from that in no-flanker trials (Fig. 4A, gray data), nor from that in congruent trials (Fig. 4A, blue data). However, when the contrast of the incongruent flanker just equaled that of the target, accuracy dropped significantly (Fig. 4A, red *, p<0.05 compared to no-flanker trials; red +, p<0.05 compared to congruent flanker trials. 2-way ANOVA followed by post-hoc comparisons with HB correction, main effect of congruency, p<0.001; main effect of contrast, p=0.021; congruency x contrast interaction, p=0.072).

**Figure 4.**
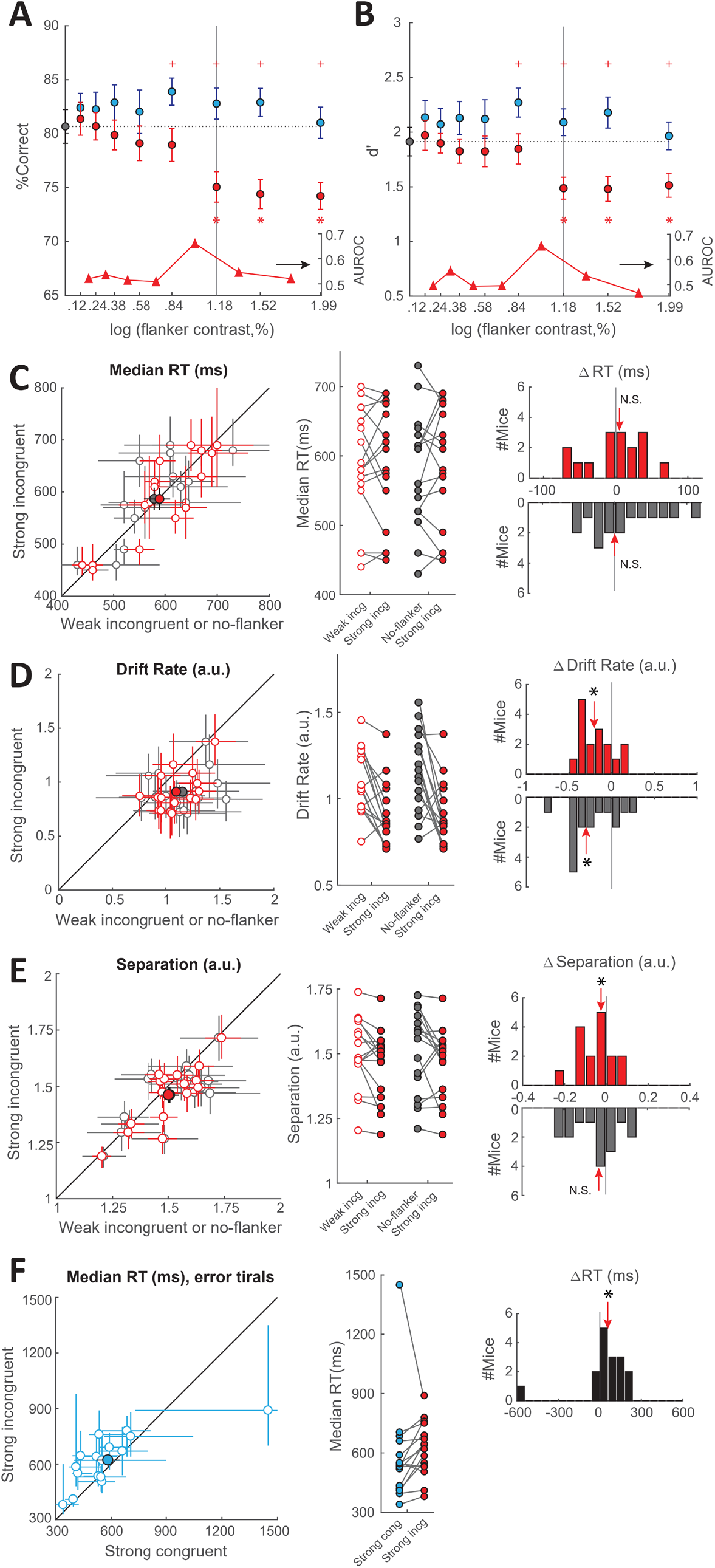
Incongruent flanker causes abrupt decrease in response accuracy and perceptual discriminability, and reduction in rate of evidence accumulation, when stronger than target. **(A-B)** Plots of response accuracy(A) and perceptual sensitivity (B) as a function of flanker contrast. Gray vertical line: fixed contrast of the target. Gray data: no flanker; blue: congruent flanker; red: incongruent flanker. +: p<0.05; incongruent trials (red) were significantly different from congruent trials (blue) of that contrast level; *: p<0.05; incongruent (red) trials were significantly different from no-flanker trials (2-way ANOVA, main effect of congruency, p<0.001, followed by post-hoc testing with correction for multiple comparisons). <u>Insets below (y-axis on rightside):</u> Ideal observer analysis quantifying area under the receiver operating characteristic (AUROC) between performance at successive pairs of contrasts in the incongruent condition, plotted at mid-point of each contrast pair. An abrupt increase in AUROC (sharp peak) was observed at flanker contrast close to that of target. Going forward, flanker contrast values 1-5 are referred to as ‘weak’ contrasts, and values 6-8, as ‘strong’ contrasts. **(C-E)** Effect of strong incongruent flanker on the median RTs (C), drift rate (D) and boundary separation (E), when compared to trials with weak incongruent flankers (red data), and trials with no flanker (gray data). Drift rate and separation obtained from drift diffusion modeling (see also Fig. S3E-G). <u>Left panel</u>:scatter plots. Each hollow circle represents individual mouse (error bars: 95% confidence interval by bootstrapping). Filled circles represent group mean ± S.E.M. <u>Mianel</u>: Comparison of value for each mouse (line) between flanker conditions. <u>Right panel</u>: Distributions of the changes (strong incongruent minus weak incongruent (red), or strong incongruent minus no-flanker (gray) trials); red arrow, median of the distribution; *: p<0.05, signed-rank and HB tests. **(F)** Comparison of the median RT in **error** trials with strong incongruent flankers versus strong congruent flankers. <u>Left panel</u>: Scatter plot. <u>Middle panel</u>: Comparison foreach mouse (line). <u>Right panel</u>: Distribution of change in median RT. *: p<0.05, signed-rank and HB tests. Seealso Figure S3.

To quantify the abruptness of this transition in performance, we employed an ideal observer analysis and computed how well responses to successive contrasts of the incongruent flanker could be discriminated (Fig. 4A, inset at bottom). This analysis revealed a large difference in the discriminability of responses to successive values of contrast, precisely when the contrast of the incongruent flanker equaled that of the target (Fig. 4A, grey vertical line).

Upon partitioning accuracy into sensitivity and criterion, we found a similar abrupt change in perceptual sensitivity as a function of flanker contrast in incongruent trials (Fig. 4B, red data and Fig. 4B, inset at bottom). There was no effect of flanker contrast on the decision criterion (Fig. S3D; 2-way ANOVA).

These results revealed that within the incongruent flanker trials, response accuracy and perceptual sensitivity were largely unaffected when flanker contrast was weaker than that of the target (‘weak contrasts’), but decreased abruptly when the flanker contrast just surpassed that of the target (‘strong contrasts’). In other words, the incongruent flanker was an effective distracter and captured attention only when it was as salient as, or more so than, the target.

Informed by these findings, we decided to examine in greater detail strong flanker trials (contrast values of #6-#8) versus weak flanker trials (with contrast values #1-#5) when the flanker was incongruent, and tested for RT differences between these two types of trials. Surprisingly, we found no difference in the median RTs (Fig. 4C, p=0.991, signed-rank test).

To investigate this effect in greater detail, we tested if there were any differences in the distributions of RTs (as opposed to just the median values), by applying the drift-diffusion modeling approach (Methods). As with diffusion modeling in the spatial probability task, we excluded outlier trials with inordinately short or long RTs. (Fig. S3E; trials with RTs between 250 ms and 1900 ms were included in the analysis, representing 95% of total trials).

We found that the drift rates were significantly lower in the strong incongruent flanker condition (Fig.4D, red data, median change= −0.207 a.u., p=0.005, signed-rank and HB tests), and the boundary separation showed a trend towards being smaller (Fig. 4E, red data; median change= −0.025 a.u., p=0.049, signed-rank and HB tests; not significant). There were no systematic differences in the other parameters (Fig. S3FG, red data; starting point, p=0.642; non-decisional constant, p=0.501, signed-rank tests). Thus, the overall absence of an effect on the RT (Fig. 4C) was the result of two competing effects: a reduction in the drift rate (consistent with distraction by the strong incongruent flanker), and trend towards reduction in ‘threshold to correct responses’ (as in the spatial probability task), but arising here from a trend towards smaller boundary separation with no difference in starting point.

The above effects also held true when the strong incongruent flanker trials were compared to the *no flanker* trials (Figs. 4C-E) and Fig. S3FG; gray data), and, as expected, also when compared to the strong *congruent* flanker trials (Fig. S3H-L).

As a final step in the analysis, we examined just the *error* trials when the flanker contrast was strong, specifically comparing the incongruent versus congruent conditions. The rationale was as follows. In the strong *congruent* flanker condition, because the flanker contains the same orientation information as the target, any attentional capture by the strong flanker is not expected to produce systematic errors. In other words, the error trials in this condition reflect errors due to ‘non-specific’ factors such as limits in learning the discrimination, failure in reporting it, and/or attending elsewhere altogether, rather than attention capture (‘distraction’) by the flanker. By contrast, the error trials in the strong *incongruent* flanker condition reflect errors due both to distraction by the flanker, as well as due to these non-specific factors. Therefore, a direct comparison of these two sets of error trials can provide an additional window, specifically, into the processes underlying attentional capture. We found that the median RTs were longer in error trials with strong incongruent flankers, compared to error trials with strong congruent flankers (Fig. 4F, median change=+57.5 ms, p=0.03, signed-rank test).

Taken together, mouse behavior in the flanker task exhibits hallmarks of attention being captured by an exogenous or bottom-up distracter (the incongruent flanker). In addition, parametric variation of the contrast of the incongruent flanker revealed an abrupt increase in the efficacy of the flanker to serve as a distracter when its salience just exceeded that of the target.

## DISCUSSION

In this study, we demonstrated that freely moving mice exhibit classic behavioral signatures of visuospatial selective attention under both endogenous as well as exogenous control. The spatially specific modulation of behavioral metrics provides evidence that the observed effects are not the consequence of general arousal, but rather of dynamic shifts in visuospatial attention. This work also departs from studies in rodents that have used variants of the 5-choice serial reaction time task (5-CSRTT) for investigations of ADHD and effects of neuromodulatory pathways (Fizet et al., 2016; Lustig et al., 2013; Robbins, 2002), because it is difficult to interpret results from those behavioral paradigms unambiguously in the context of selective visuospatial attention. Complementing a recent study in head-fixed mice (Wang and Krauzlis, 2018), these results help establish a reliable, primate-inspired behavioral foundation in mice to study selective visuospatial attention and to investigate the underlying neural circuit mechanisms.

### Studying selective attention in freely behaving animals

Notably, our results establish a parameterized approach for studying selective visuospatial attention in unrestrained animals – the first such approach to the best of our knowledge. Head-fixed animals, with the advantages they afford for tightly controlled experiments, have been used extensively in neuroscience research (and exclusively in research on selective visuospatial attention). However, recent work has shown explicitly that there are significant differences in neural representations even between stationary versus locomotory behavioral states (Cardin and Schmidt, 2004; Cohen and Castro-Alamancos, 2010; Fu et al., 2014; Keller et al., 2012) In addition, compared to head-fixed animals that are capable of tethered motion (walking, running or flying), unrestrained animals receive a full complement of proprioceptive and vestibular cues that play an important part in the control of natural behavior (Aghajan et al., 2015; Chen et al., 2013; Rancz et al., 2015; Ravassard et al., 2013). The ability to characterize complex behavior under such conditions, therefore, affords the power to obtain a richer picture of the neural underpinnings of that behavior. This holds true especially for selective visuospatial attention, a function that animals employ heavily when interacting freely with their environments. Nonetheless, a key challenge that any experimental paradigms involving free behavior must address is the potential reduction in the degree of experimental control and increase in variability. Here, we demonstrate that freely behaving mice exhibit systematic changes in behavioral performance of similar magnitudes as those reported in head-fixed mice (Wang and Krauzlis, 2018). Therefore, these results confirm the reliability of our approach for studying visuospatial selective attention in fully unrestrained animals. Together with the emergence of tools for genetically targeted measurement and manipulation of neural activity in unrestrained mice, this study sets the stage for dissection of the neural circuitry underlying spatial attention control.

### General features of behavioral tasks in this study

Our tasks involved presenting stimuli at two possible locations along the vertical axis: one in the upper periphery and the other closer to the center. The asymmetry in visual perceptual performance that we find between these two locations is similar to that reported in humans (Carrasco et al., 1995; Carrasco et al., 2004; Carrasco et al., 2001).

The range of performance observed even in the relatively simple, single-stimulus discrimination task had a median of 84.3% (Fig. S1B), which is low compared to the levels of performance (upper 90s) on similar tasks in primates (Vazquez et al., 2000; Vogels and Orban, 1990). This difference is accounted for by a combination of the lower visual acuity of mice compared to primates, and our use of ‘minimal’ stimulation (small stimuli). Indeed, with full field grating stimuli, mice can perform very well (accuracy in the upper 90s) (Andermann et al., 2010; Long et al., 2015). We chose to use small stimuli because this was necessary to test for spatial competition and spatially specific effects. Notably, the lower range of accuracy did not pose any obvious confounds because we were still able to detect changes in performance due to attention, effects of the same kind as those reported in human and primate attention tasks.

Our tasks were designed as 2-alternative forced choice discriminations, as opposed to yes-no detection tasks, thereby minimizing overall response bias (Gomez et al., 2007), and notably, allowing the quantification of RTs for trials corresponding to both choices (i.e., trials with vertical as well as horizontal targets, in our case).

A key goal of this study was to develop tasks that dissociated the locus of sensory stimulus/attention from that of animals’ reports, so as to be unambiguously able to attribute any observed changes in behavior to motor vs. sensory or cognitive sources. This goal was achieved by orthogonalizing the axes of stimulus locations and nose-touch locations, and training all mice at the outset on a fixed response-association rule (Fig. 1). As a consequence of this design, we were able to rule out alternative explanations to attention for the results (Figs 1–2 and S1; see also next subsection). An added advantage of this design is that it allows for the behavioral paradigms developed here to be used directly in future neural recording and perturbation experiments.

### Aspects of the spatial probability task

Our spatial probability task is based on those previously used in human studies (Druker and Anderson, 2010; Geng and Behrmann, 2005; Vincent, 2011). Specifically, in our task, spatial expectation was manipulated by blocked changes in the probability of target occurrence at the upper versus lower locations (90:10 vs. 50:50). As a result, in the 90u-10 blocks, the number of trials with the target at the lower location was necessarily much smaller than the number of trials at the upper location (1:9), accounting for the increased variability in the data obtained at the lower location (larger error bars of purple data; Figs. 1DE, S1KMN, 2, S2C-F).

In general, the magnitude of effects observed in the lower location was larger than of those at the upper location (for instance, the reduction in accuracy and d’ at lower location were larger than the corresponding increases at the upper location; Fig. 1DE). This difference potentially reflects limits of perceptual behavior at the upper location, consistent with the observation of poor baseline performance at the upper location (Fig. S1H). Regardless, behavioral metrics revealed significant, space-specific effects on performance.

Notably, these effects were best explained by endogenous expectation-driven shifts of spatial attention rather than by any of the five possible alternatives. First, the space-specific nature of the observed effects ruled out general arousal as a potential explanation for them. Second, the results could not be accounted for by potential differences in sensory input because we found no systematic differences in sensory input between block-types at either target location (Fig. S1M). Third, they could not be accounted for by low-level sensory after-effects arising from sequential trial structure or repetition priming (Hillstrom, 2000; Kristjansson et al., 2002; Maljkovic and Nakayama, 1996; Quinlan and Hill, 1999). This is because, by virtue of the 2-AFC design of the task together with the randomization of the identity and location of target presentation, the probability of a vertical (horizontal) stimulus immediately following a vertical (horizontal) stimulus when both stimuli were presented at the upper location (sequential trials), was not different between the 90u-10 and 50-50 conditions. Moreover, the probability of a correct response on the second of two successive trials in which the same target was presented at the upper location, was also not different between the 90-10 and 50-50 conditions (Fig. S1Q). Fourth, they could not be explained by differences in motor biases, because our task design required operant responses at locations distinct from the target locations. Indeed, we found no differences in the left-right responding of the animals between the block-types (Fig. S1N). Fifth and finally, the inter-trial intervals were long (median > 15 s in both block-types), with the mice traveling to the reward port, consuming reward, and returning to the touchscreen to initiate the next trial, during this period. Consequently, the observed effects are not explained by mnemonic motor strategies, such as postural mediation (Dudchenko, 2004), to mark the location of higher probability. Our results are, therefore, best explained by the mice inferring and holding information about the spatial probabilities in working memory, in other words, by the action of an endogenous influence (Goldman-Rakic, 1995; Knudsen and Knudsen, 1996; Liu et al., 2014; Rikhye et al., 2018).

Our task involved the manipulation of spatial probability to effect endogenous control of spatial attention. Human studies have shown that probability manipulation can itself serve as a spatial attentional cue (Druker and Anderson, 2010; Geng and Behrmann, 2005). Indeed, direct comparison of results between trials with an explicit spatial cue versus with a probability manipulation revealed no differences from a Bayesian observer’s perspective (Vincent, 2011). In this context, in the 90u-10 blocks of the spatial expectation, trials with the target at the upper location may be considered as trials with a ‘valid’ cue (90% of trials), whereas trials with the target at the lower location may be considered as trials with an ‘invalid’ cue (10% of trials). In primate studies, it is standard practice to treat validly cued trials as the ‘attend towards’ condition, and the invalidly cued trials as the ‘attend away’ condition, and to characterize the effect of attention as the difference in performance between the two ‘attention’ conditions. We performed this comparison as well, by computing the differences in performance between the upper and lower locations in the 90u-10 blocks (Fig. S1OP; maroon data), relative to those in baseline (the 50-50 block; Fig. S1OP; teal data). We found an improvement of 6.5% in accuracy (difference between the medians) and 0.57 s.d. in d’. In other words, these are the net improvements in the ‘attend towards’ over ‘attend away’ conditions.

Whereas the arguments described above are based on the finding that probability manipulation can itself serve as a spatial attentional cue (Druker and Anderson, 2010; Geng and Behrmann, 2005; Vincent, 2011), an alternate approach used to great effect in primates to study endogenous control of attention involves the use of an explicit spatial cue that predicts the target’s location, with the position of the cue randomized from trial to trial (Bashinski and Bacharach, 1980; Posner et al., 1980). Such a task has not been reported in mice thus far. Although a recent study in head-fixed mice utilized a spatial cue (Wang and Krauzlis, 2018), because its location was varied in a blocked fashion rather than randomly between successive trials, it is difficult to rule out that the reported effects are due to blocking-dependent changes in spatial probability, as in our task. In this context, the ability of our mice to infer spatial probabilities and shift attention appropriately even in the *absence* of an explicit spatial cue, and in a freely behaving condition, are significant.

In this task, and more generally, in this study, we did not measure the gaze directions, i.e., eye-in-orbit and head-in-space positions, of mice during task performance. However, this was not necessary for the unambiguous interpretation of our results because our goal was the behavioral demonstration of visuospatial selective attention, rather than demonstration, specifically, of overt versus covert versions of it.

### Drift diffusion modeling of RTs

In both the tasks described here, the use of drift diffusion modeling to analyze the distributions of RTs yielded additional insights beyond characterization of just their median values, with effects on the drift rates providing strong support for shifts of visuospatial attention. Additionally, in the spatial probability task, the observed reduction of the ‘threshold to correct responses’ at the lower probability target location, was able to explain the seemingly puzzling speeding-up of reaction time at that location.

In this context, literature suggests that when the diffusion model is set up with the two ‘choices’ being correct versus incorrect responses (as opposed to responses to vertical versus horizontal gratings, for instance), changes in thresholds to correct (or incorrect) responses are difficult to interpret, and recommends fixing the starting point at halfway along the boundary separation (Voss et al., 2015). Our use of a variable starting point, and conclusions regarding changes to ‘threshold to correct responses’, although seemingly at odds with this viewpoint, are, in fact, lawful. In our task, information about spatial probabilities causes changes in response accuracy. Therefore, the experimental block (90u-10 vs. 50-50) can intrinsically alter the likelihood of producing correct responses, indicating that ‘threshold to correct responses’ is a valid quantity in the context of our task design.

Taken together, then, the effects on drift rates and thresholds revealed by diffusion modeling enabled the generation of detailed insights into seemingly puzzling changes in median RT values.

### Aspects of the flanker task

Our touchscreen-based flanker task is based on the classic flanker task of attention in humans (Eriksen and Eriksen, 1974; Fan et al., 2002) in which comparisons are made between the incongruent and congruent conditions (and in some cases, also between the incongruent and single-stimulus conditions). The effects we report here are consistent with that literature (Eriksen and Eriksen, 1974; Fan et al., 2002). Notably, our task goes a step further and parameterizes the salience of the flanker.

Our parameterized design highlights an important aspect of the flanker task that is not always emphasized. Although the task demonstrates exogenous capture of attention, both exogenous as well as endogenous influences are major factors in this task. It explicitly requires mice to pay attention to the fixed location of the (reward-yielding) target, resulting in attention being directed to the target in an endogenously driven manner. As revealed by our results, it is only when the flanker becomes sufficiently salient that it is able to overcome the endogenously highlighted target to capture attention (Fig. 4AB, data points 6-8 at right end). In trials in which the flanker contrast is less salient, there is no observable difference in performance from either the single target condition or the congruent flanker condition, indicating the inability of the flanker to overcome the endogenous influence to direct attention to the central target. Thus, each trial of this task involves constant competition between purely exogenous influences due to the flanker stimulus, and endogenous plus exogenous influences towards the target, with the former winning out when the flanker is strong, resulting in an abrupt drop in response accuracy and d’.

As a result, our parameterized task design is an excellent substrate for future investigations into the neural signatures of priority-dependent target selection and distracter suppression in the oculomotor pathway, with the observed switch-like change in behavioral metrics suggesting that the task may additionally be well-suited to reveal whether these neural signatures of attentional selection are explicitly categorical, as hypothesized recently (Mysore et al., 2011; Mysore and Knudsen, 2014).

## ACKNOWLEDGEMENTS

This work was supported in part by funding from NIH R03 HD093995. We are grateful to Drs. Howard Egeth and Daniel O’Connor for feedback on the manuscript, Drs. Veit Stuphorn and Patricia Janak for comments on the research, and James Garmon for help with fabrication of custom equipment.

## AUTHOR CONTRIBUTIONS

SPM and WKY designed the research and wrote the paper, and WKY performed the experiments and analyzed the data.

## METHODS

### Animals

All mice were C57Bl6/J strain purchased from the Jackson Laboratory. Upon arrival, mice were housed in a colony where temperature (∼75F) and humidity (∼55%) were controlled on a 12:12h light : dark cycle. At least one week of acclimation period was allowed with food and water *ad libitum* before water restriction was initiated. Experiments were all carried out in the light phase. All procedures followed the NIH guidelines and were approved by the Johns Hopkins University Animal Care and Use Committee (ACUC).

### Water restriction

Mice were water-restricted following protocols described by Guo et al. (2014) with a few modifications. Briefly, mice were individually housed, and administered 1mL water per day to taper their body weight down to 80-85% of each animal’s baseline, over the course of 5-7 days. During behavioral training/testing, the primary source of water for mice was as a reinforcer for correct performance: 10 µL of water was provided for every correct response.

### Apparatus

Behavioral training and testing were performed in soundproof operant chambers equipped with a touchscreen (Med Associates Inc.), a custom-built reward port (fluid well), infrared video cameras, a house light and a magazine light above the reward port. The reward port was located at the opposite wall of the chamber relative to the touchscreen (Fig. 1A). Two custom modifications were introduced that limited the area of the touchscreen available for exploration by the freely behaving mice, thereby minimizing false-alarm triggers due to accidental touches. First, mice were placed within a clear plexiglass tube that ran from the touchscreen to the reward port. The diameter of the tube (5 cm) was large enough to allow mice to run back and forth from the touchscreen to the reward port, to groom and to behave naturally. Second, a thin plexiglass mask (3 mm thickness) was placed 3 mm in front of the touchscreen with three holes corresponding to the locations at which the mouse was allowed interact with the screen by a nose-touch (Fig. 1A). The holes, each 1 cm in diameter, were drilled in the mask in an inverted triangle configuration: ‘left’ and ‘right’ holes were placed 3cm apart (center-to-center) along the base of the triangle, and a ‘central’ hole, at the apex of the triangle, was 1.5 cm below the midpoint of the base (Fig 1A). All experimental procedures were executed using control software (K-limbic, Med-Associates).

### Visual stimuli

Visual stimuli (bright objects on a dark background; background luminance = 1.32 cd/m^2^) were generated using MATLAB (Mathworks) and imported into the K-Limbic system as jpeg images. A small cross (60×60 pixels; luminance = 130 cd/m^2^) was presented in the central hole and had to be touched to initiate each trial. The experimental stimuli were oriented gratings (horizontal or vertical orientation; 60×60 pixels), generated using a square wave (24 pixels/cycle; within the range of spatial frequencies shown to be effective for mice; (Histed et al., 2012)). The dark phase of the cycle was black (luminance = 1.32 cd/m^2^; same as the background), and the bright phase was varied between 1.73 cd/m^2^ and 130 cd/m^2^ to control its contrast (flanker task, see below). The stimuli subtended 25 visual degrees at a viewing distance of 2 cm from the screen.

### Experimental procedure and behavioral training

Each mouse was run for one 30 min behavioral session per day, with each session yielding 80-180 trials. Each behavioral session began with a 10 s acclimation period, during which mice were allowed to explore the environment with the lights on and to retrieve a bolus (10 µL) of ‘free’ water at the reward port. Following this, lights shut off and the zeroing cross to start the first trial appeared on the screen. The cross flashed once every 10s until touched, and the flash was accompanied by a short beep of 600 Hz for 30 ms, to induce the mouse to approach and begin the trial. Upon trial initiation, the cross vanished, and the visual stimulus (or stimuli) were immediately presented for a duration of 1-3s depending on the task (see below).

Mice were trained to report the information contained in the target grating, namely, its orientation, by nose-touching within the correct response hole (vertical target grating → nose-touch in left response hole; horizontal target grating → nose-touch in right response hole). A correct response triggered a tone (600 Hz, 1 sec), the turning on of the magazine light above the reward port, and the delivery of 10 microliters of water at the reward port. Mice turned away from the screen, ran to the liquid well, consumed the reward, and ran back to face the touchscreen in order to begin the next trial. Mouse head entry into the reward port was detected by an infrared sensor which caused the magazine light to turn off, and the zeroing cross (for the next trial) to be presented on the touchscreen. An incorrect response triggered the turning on of both the house light and the magazine light for 5-s as a punishment/timeout; the next trial could not be initiated until the end of timeout. A failure to respond within 3s of stimulus presentation resulted in the stimulus vanishing and the zeroing cross being presented immediately (without a timeout penalty) for initiation of the next trial. Well-trained animals failed to respond on fewer than 5% of the total number of trials, and there were no systematic differences in the proportion of such missed trials between different conditions.

Within each daily 30-minute behavioral session, mice consumed approximately 1mL of water. If a mouse failed to collect enough water from the behavioral session, they were provided with a water supplement using a small plastic dish in their home cage. The specific amount of supplement was customized depending on individual animal’s body weight, the training phase it was in, and the motivational drive observed during the experiment.

### Single-stimulus discrimination task

Upon trial initiation, a single, full contrast, grating stimulus (‘target’, 60×60 pixels, 25°, 2.5 cycle) was presented above the central hole, aligned along the elevation with the left and right holes. The stimulus was presented for a duration of 3s, and mice were required to report its orientation with the appropriate nose-touch.

### Spatial probability task

Upon trial initiation, a single, full contrast, grating stimulus (‘target’, 60×60 pixels, 25°, 2.5 cycle, 2s) was presented at one of two possible locations: an upper location (the center of the grating was 90 pixels or 37.5° above the center of the central hole), and a lower location (30 pixels or 12.5° above the central hole). The stimulus was presented for a duration of 2s, and mice were required to report its orientation with the appropriate nose-touch. Trials were run in blocks of two kinds, with different probabilities of target occurrence at the upper (and lower) locations; blocks were interleaved pseudorandomly across days. In the ‘50- 50’ blocks, the probability that the target would appear at the upper location on any trial was 50% (and at the lower location, also 50%). In the ‘90u-10’ blocks, the probability, on any trial, that the target would appear at the upper location was 90% (and at the lower location, 10%). This design allowed us to test if spatial expectation altered behavioral performance. To train mice on this spatial probability task, they were first trained on the single stimulus discrimination task (with the target always presented midway between the lower and upper locations), following which the two possible target locations with corresponding spatial probabilities were introduced.

### Flanker task

Upon trial initiation, either one stimulus (‘target’, 60×60 pixels, 25°, 2.5 cycle, 25%) was presented at the lower location, or two stimuli were presented simultaneously, with the target at the lower location and a second ‘flanker’ at the upper location. Flankers were of the same size and spatial frequency as the target, but of contrast in 8 different levels: 1.31%, 1.73%, 2.42%, 3.79%, 6.84%, 15.2%, 33.4%, 98.5%. The orientation of the flanker was either identical to that of the target (‘congruent trial’) or orthogonal to that of the target (‘incongruent trial’). The stimulus (stimuli) was (were) presented for a duration of 1s, and mice were required to report orientation of the target grating with the appropriate nose-touch. All types of trials (no flanker, congruent, incongruent) and flanker contrasts were interleaved randomly within each daily session. To train mice on this flanker task, they were first trained on the single stimulus discrimination task (with the target always at the lower location), following which, a flanker was introduced at the upper location with progressively increasing contrast over training days.

### Subject inclusion/exclusion

A total of 25 mice were used in this study. All 25 were trained on the single stimulus discrimination task, and of these, n=20 mice satisfied the inclusion criterion: % correct > 70% (yielding the data shown in Fig. S1B-G). All 25 mice were also trained on the spatial probability task, and of these, n=17 mice satisfied the inclusion criterion for the spatial probability task: overall % correct in the (baseline) 50-50 condition > 70% (data shown in Figs 1, S1I, K-Q and 2). A total of 18 mice were trained on the flanker task, and of these, n=16 mice satisfied the inclusion criterion for the flanker task: overall % correct across all flanker contrasts >70% (data shown in Figs 3 and 4). Between experiments, mice were well rested for at least few weeks with food and water *ad libitum*, before the next water restriction was initiated.

### Behavioral measurements

Response accuracy (% correct) was calculated as the number of correct trials divided by the total number of trials responded (correct plus incorrect). Reaction time (RT) was defined as the time between the start of stimulus presentation and response nose-touch, both detected by the touchscreen.

### Signal detection analysis (sensitivity and criterion)

In the framework of signal detection theory, we assigned the correct vertical trials as ‘hits’, incorrect vertical trials as ‘misses’, correct horizontal trials as ‘correct rejections’ and incorrect horizontal trials as ‘false alarms’, and calculated the perceptual sensitivity (d’) and criterion (c) accordingly (Stanislaw and Todorov, 1999). Because of the inherent symmetry in 2-AFC tasks, this calculation was independent of which grating orientation – vertical or horizontal – was assigned as ‘signal’ and which as ‘noise’. Consequently, a positive value of c caused poor performance just as much as the corresponding negative value, and therefore, we quantified the absolute value of c (|c|) as the relevant metric of decision criterion (Figs. 1, 3 and 4).

### Trial inclusion/exclusion

Towards the end of a behavioral session, when mice had received a sizeable proportion of their daily water intake, they were observed to become less engaged in the task. This was reflected in their behavioral metrics: they tended to wait longer to initiate the next trial, and their performance deteriorated. To avoid confounds due to loss of motivation towards the end of sessions, we developed an unbiased procedure to identify and exclude such trials. To this end, we pooled data across all mice and all sessions, treating them as coming from one session of a single ‘mouse’. We then binned the data by trial number within the session, computed the discrimination performance in each bin (% correct), and plotted it as a function of trial number within session (Figs. S1DI, S3A). Additionally, we used a bootstrapping method to compute the 95% confidence interval for this value. As expected, we found that the performance became highly variable and dropped towards chance for trials towards the end of the session (Figs. S1DI, S3A). Using this data, we developed the following exclusion criterion: Trials q and above were dropped if the qth trial was the first trial at which *at least one* of the following two conditions was satisfied: (a) the performance was statistically indistinguishable from chance on the qth trial and for the majority (3/5) of the next 5 trials (including the qth), (b) the number of observations in qth trial was below 25% of the maximum possible number of observations for each trial (mice*sessions), thereby signaling substantially reduced statistical power available to reliably compare performance to chance.

For the single stimulus discrimination task, q was determined to be the 122nd trial using pooled data from all the sessions of the 20 mice (Fig. S1D). Thus, trials 122 and above were dropped from each session. This resulted in the exclusion of 3.7% of the trials (229 out of 6176 trials).

For the spatial probability task, q was determined to be the 141st trial using pooled data from all the 50-50 sessions (irrespective of the stimulus location) of the 17 mice (Fig. S1I). Thus, trials 141 and above were dropped from each 50-50 session of the spatial probability task. This resulted in the exclusion of 5.7% of the trials (781 out of 13772 trials). This same value of q was used to drop trials from the 90-10 sessions to ensure unbiased treatment of both conditions; this resulted in the exclusion of 4.10% of the trials (656 out of 15522 trials).

For the flanker task, q was determined to be the 131st trial using pooled data from all sessions (irrespective of the trial type and flanker contrast) of the 16 mice (Fig. S3A). Thus, trials 131 and above were dropped from each session of the flanker task. This resulted in the exclusion of 2.53% of the trials (834 out of 32959 trials).

### Drift diffusion modeling of RT distributions

To shed light on potential mechanisms underlying observed RT distributions, we applied the drift-diffusion model to our RT data (Voss et al., 2015). This model hypothesizes that a subject (‘decision maker’) collects information from the sensory stimulus via sequential sampling, causing sensory evidence to accrue for or against a particular option (usually binary) during the viewing of the stimulus. A decision is said to be made when the accumulating evidence reaches an (abstract) internal threshold of the subject. This process of evidence accumulation, together with the processes of sensory encoding and motor execution, as well as threshold crossing, are said to determine the RT observed on each trial.

We used a standard version of the model that consists of four independent variables (Ratcliff, 1978; Ratcliff et al., 2016; Voss et al., 2013): (1) the drift rate, which represents how fast sensory evidence accrues; (2) the boundary separation, which represents the bar that subject sets for the decision to be made; (3) the starting point, which represents the (prior) biases the subject might have favoring one versus the other option (=0.5 when unbiased); and a (4) non-decisional constant, which accounts for the time spent in sensory encoding and motor execution. In the case of our tasks, there was no reason for the drift rate to be different between vertical versus horizontal gratings, and therefore, we merged both type of trials (trials with a horizontal target grating and trials with a vertical target grating). We treated ‘correct’ response and ‘incorrect’ response as the two binary options, and fit the diffusion model to the RT distributions of correct versus incorrect trials using the fast-DM-30 toolbox with the maximum likelihood option to gain estimates of those four parameters for each individual mouse (Voss et al., 2015).

To obtain accurate estimates of model parameters, it is important to drop trials with outlier values of RTs (too fast or too slow trials), as per the established approach in drift-diffusion modeling (Voss et al., 2015). We developed an unbiased procedure to identify and exclude inordinately fast or slow trials. We reasoned that on trials with RTs that are so short as to not allow mice sufficient time to accumulate sensory evidence, performance would be consistently poor because mice would be forced to guess. Similarly, on trials with RTs that are so long (far exceeding stimulus offset) as to extinguish the trace of sensory evidence from their short-term memory (Ratcliff and Rouder, 2000; Ratcliff et al., 2016; Smith and Ratcliff, 2009), performance would be consistently poor because animals would be forced to guess. To apply this heuristic, we pooled RT data across all mice and all sessions, treating them as coming from one session of a single ‘mouse’. We then binned this RT distribution into 50 ms bins and for each bin, computed the response accuracy and the 95% confidence intervals (using a bootstrapping method). We then identified short and long RT bins for which the response accuracy was statistically indistinguishable from chance (Fig. S2B, S3E).

For the spatial probability task, using pooled data from all 50-50 sessions of the 17 mice (irrespective of the stimulus location), we determined that trials with RTs shorter than 300 ms or longer than 2550 ms were outliers (Fig. S2B). This resulted in the exclusion of 6.3% of the trials (815 out of 12991 trials). The same RT exclusion range was used for data from the 90u-10 sessions to ensure unbiased treatment of both conditions. This resulted in the exclusion of 9% of the trials (1335 out of 14866 trials).

For the flanker task, using pooled data from all sessions of the 16 mice (irrespective of trial type and flanker contrast), we determined that trials with RTs shorter than 250 ms or longer than 1900 ms were outliers (Fig. S3E). This resulted in the exclusion of 5.06% of the trials (1625 out of 32125 trials).

### Statistical tests

All analyses and statistical tests were performed in MATLAB. *<u>Cross-condition comparisons (Figs. 1–4).</u>* For each behavioral metric (% correct, RT, etc.), the non-parametric Wilcoxon signed-rank test was used to test if the median of the distribution of the change in the metric between conditions (e.g., 90u-10 vs. 50- 50, or incongruent vs. congruent) was different from zero, followed by the Holm-Bonferroni test for multiple comparisons across stimulus locations (spatial probability task) or flanker conditions (flanker task). The application of this statistical procedure is referred to in an abbreviated fashion as ‘signed-rank and HB tests’ in the text. *<u>Correlation analysis (Figs. 1FG</u>*, *<u>3F).</u>* The Pearson correlation coefficient was calculated for paired data, and p-value was computed using Student’s t-test after transforming the correlation into a t-statistic having n-2 degrees of freedom. *<u>Flanker contrast-dependent analysis (Fig. 4AB)</u>*. 2-way ANOVA was first used to examine the effect of flanker congruency, and the effect of flanker contrast. Post-hoc paired comparisons were then performed using Student’s t-test in selected pairs, followed by the Holm-Bonferroni test (HB test) for multiple comparisons.

### Code and data availability

Software code and the data that support the findings of this study are available from the corresponding author upon reasonable request.

## Supplemental videos

**Video S1**. Video snippet showing mouse performing the 50-50 block of the spatial probability task.

**Video S2.** Video snippet showing mouse performing the flanker task.

