## Supplemental Figures for "Endogenous and exogenous control of visuospatial selective attention in freely behaving mice"

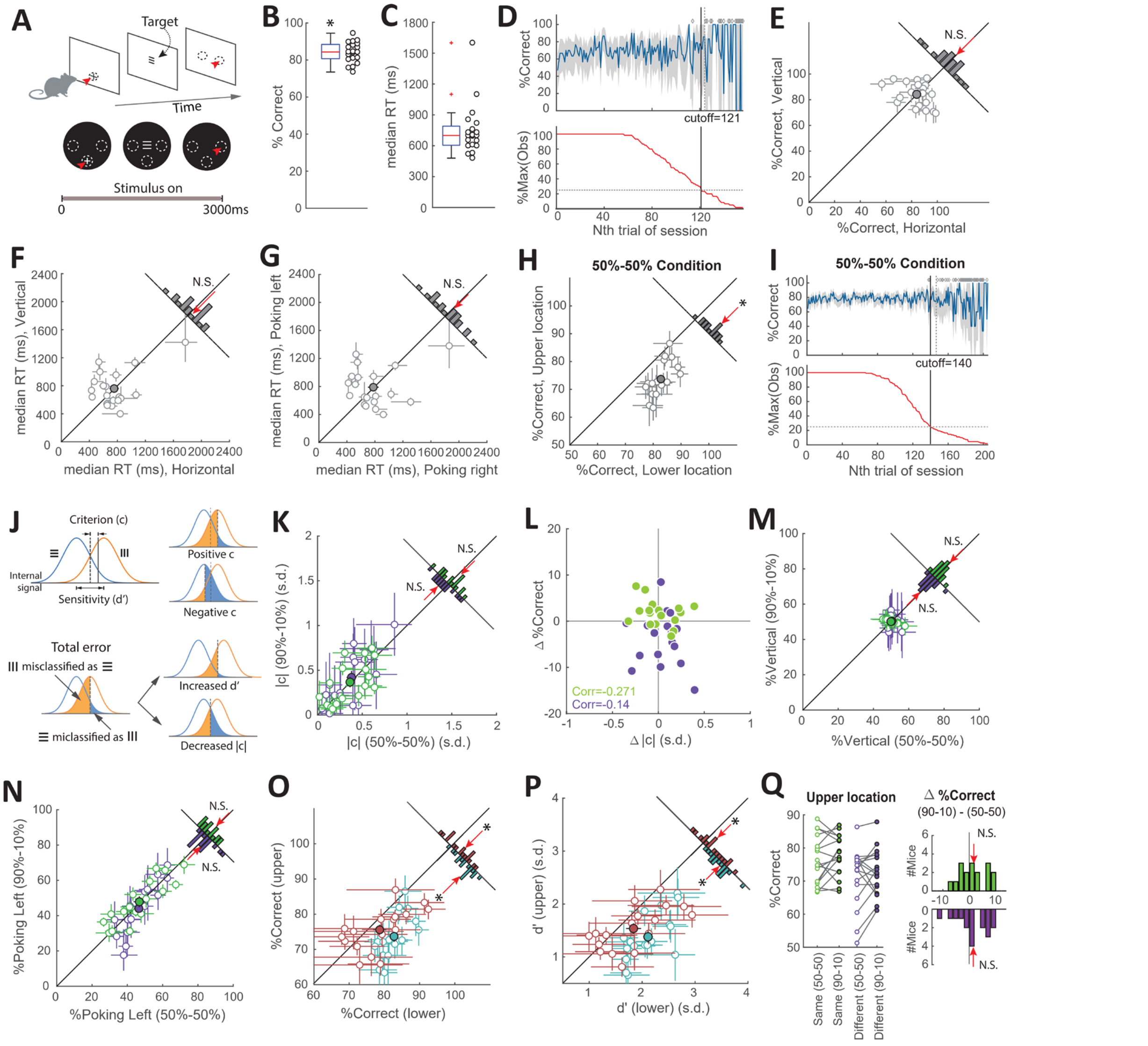


**Figure S1. Related to Figure 1. Single-stimulus discrimination task and spatial probability task: trial exclusion, location asymmetry, signal detection theory (SDT), and lack of sensory or motor biases.**

**(A-G)** Single-stimulus visual discrimination task: Freely behaving mice learn feature-response association. **(A)** Schematic of task design. _Top row:_ Trials began with a nose-touch on a zeroing-cross presented within the lower central hole. A single oriented grating ('target') was presented after trial initiation (size= 25$^{o}$, 60 pixels x 60 pixels; duration = 3s, contrast=98.5%; Methods). Mice were rewarded if they responded to the target per the following rule: vertical target grating → nose-touch in left response port; horizontal target grating → nose-touch in right response port. Dashed ovals: response holes; the 'zeroing' hole is shown only in the first stage, and the two response holes are shown only in the third stage for clarity. Red arrowheads denote nose-touches. _Bottom-row:_ Screen-shots of display at different stages in a trial. **(B)** Response accuracy; median = 84.3%. **(C)** Median reaction time (RT); median value = 700 ms. Each circle denotes data from one mouse, calculated after pooling its trials from all behavioral sessions (Methods). Data are from n=20 mice. Box-plot summarizes the distribution; *: statistically significant from chance level (signed-rank test); '+' denotes outliers. **(D)** Identification of trials (late in sessions) that corresponded to animals being poorly engaged in the task (Methods). _Top panel_: Timecourse of overall response accuracy across mice as a function of trial number within sessions. Accuracy obtained from trials pooled across all mice and sessions, and computed as a function of trial number within session (blue; Methods). Grey shading: bootstrapped estimates of the 95% confidence interval of the accuracy (gray; Methods). Open diamonds: accuracy not significantly different from chance for corresponding trial. Dashed vertical line: first trial at which the accuracy was not different from chance (50%), and stayed indistinguishable from chance for 3/5 of the next 5 trials

(Methods). Data show increased variability and worse performance towards the end of sessions. *Bottom panel*: Number of actual observations across mice for each trial number, as a percentage of the maximal number of possible observations (mice * sessions), plotted as a function of trial number within session (red). Solid vertical line: first trial at which the number of observations drops below 25%. Data show drop in the number of observations available to reliably assess performance towards the end of sessions. Based on these data, all trials above 121 (black vertical line) of each behavioral session of this task were dropped from analysis (Methods). Results in panels S1BC, S1E-G are based on data from trials 1-121 from each behavioral session. **(E-F)** Scatter plot of response accuracy (E) and median reaction time (RT; F) when the target was a horizontal grating versus when it was a vertical grating. **(G)** Scatter plot of median RT to nose-touches to right versus left response holes.
**(H-P)** Spatial probability task. **(H)** Scatter plot comparing response accuracy (%-correct) at the upper location versus the lower location in the (baseline) 50-50 condition. *Inset:* Distribution of difference in performance between the upper and lower locations. Red arrow: median; * $p<0.05$; signed-rank test. **(I)** Identification of trials (late in sessions) that corresponded to animals being poorly engaged in the task (Methods; conventions as in D, above). *Top panel*: Overall response accuracy versus trial number. Accuracy obtained from trials pooled across all mice and locations from 50-50 sessions, and computed as a function of trial number within session (blue; Methods). *Bottom panel*: Number of actual observations across mice for each trial number, as a percentage of the maximal number of possible observations (mice * sessions), plotted as a function of trial number within session (red). Based on these data, all trials above 140 of each behavioral session of this task were dropped from analysis (Methods). Results in Figures 1 and 2 are based on data from trials 1-140 from each behavioral session.
**(J)** Schematic of the signal detection theory (SDT) analysis. *Upper row; left*: SDT hypothesizes that the internal representation of vertical and horizontal stimuli can be reduced (projected) to a one-dimensional decision axis, on which they form two overlapping distributions (due to noise). A decision is made based on a criterion set by each individual animal: a stimulus whose representation falls above (or below) the criterion is judged as vertical (or horizontal), producing the appropriate behavioral response. The decision criterion (c) here is quantified as the amount of deviation from a neutral (unbiased) value (where c=0). *Upper row; right*: Because of the symmetry of 2-AFC task design, deviations of the criterion to the left (i.e., negative, favoring vertical stimulus) or to the right (i.e., positive, favoring horizontal stimulus) of the neutral value produces the same effect on overall response accuracy (despite potential differences in accuracy in vertical versus horizontal trials). For this reason, we used the absolute value of c (|c|) to examine the effect of criterion change on response accuracy. *Lower row*: Based on theory, improved response accuracy can result from (1) increased d': when the two distributions become further separated; or (2) decreased |c|: when the decision criterion becomes less biased. **(K)** Scatter plot comparing the criterion in the 90u-10 versus 50-50 condition at the upper (green data) and lower (purple data) locations. **(L)** Change in response accuracy plotted against change in criterion at the upper (green) and lower (purple) locations; they were not correlated (Pearson's correlation; $p>0.05$). **(M)** Scatter plot comparing percentage of vertical stimulus presentations between the two conditions at the upper (green) and lower (purple) locations. **(N)** Scatter plot comparing percentage of left nose-touches between the two conditions at the upper (green) and lower (purple) locations. No significant change in both cases ($p>0.05$; signed-rank test), indicating that differences in sensory input or motor biases cannot account for the effects in Fig. 1. **(O)** Scatter plot comparing response accuracy (% correct) at the upper versus lower locations in the 50-50 condition (teal) and 90u-10 condition (maroon). Inset shows distributions of changes in accuracy between the upper and lower locations for the two conditions. Red arrows: medians; *: $p<0.05$, signed-rank and HB tests. **(P)** Scatter plot comparing perceptual sensitivity accuracy (d') at the upper versus lower locations in the 50-50 condition (teal) and 90u-10 condition (maroon). Conventions as in O. **(Q)** Examination of sequential effects at the upper location. *Left panel:* For successive trials in which the target was presented at the upper location, plot of probability of a correct response on the second trial when the target was the same in both trials (green data), and when the target was different in the two trials (purple data). Each line denotes data from one mouse. *Right panel:* Distribution of change in probabilities (across n=17 mice) between the 90u-10 and 50-50 conditions; colors as in left panel.

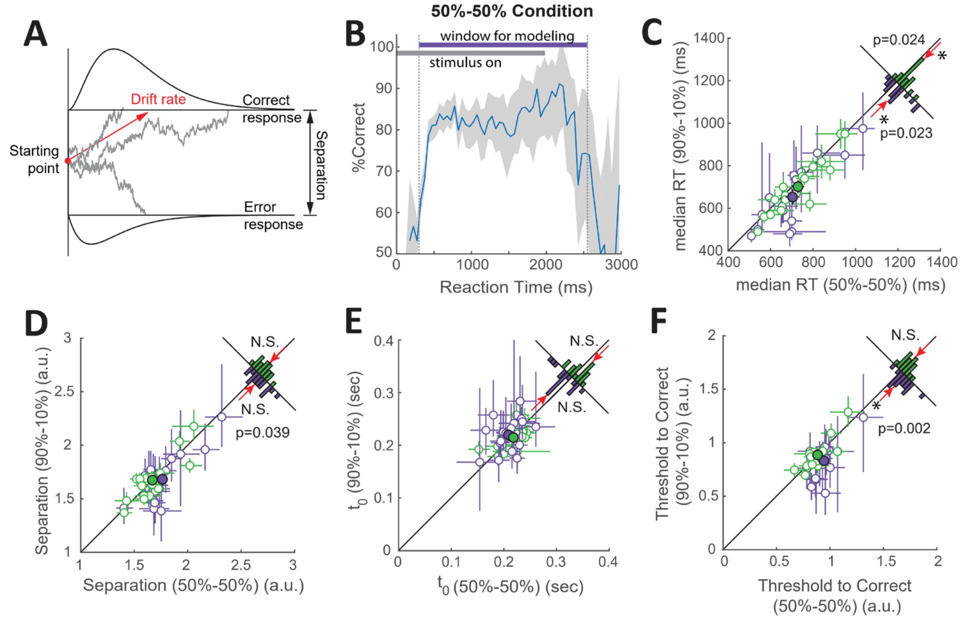


**Figure S2. Related to Figure 2. Spatial probability task: Drift diffusion modeling**
**(A)** Schematic diagram of the two-choice drift diffusion model. The model simulates a decision process from sensory stimulus presentation to the point of behavioral report, and attempts to account for the full distribution of observed RTs. It posits that upon stimulus presentation, sensory evidence flows in, causing a (hypothetical) decision variable to 'drift' either upwards (towards one choice boundary) or downwards (towards the other) depending on which choice the incoming evidence favors. Under uncertainty, the decision variable drifts in a stochastic (zig-zag) manner as sensory evidence accumulates, eventually crossing one of the decision boundaries and triggering the corresponding behavioral response. Here, we adopted a standard version of the model with four parameters: (i) drift rate, or the average rate of evidence accumulation, whose sign could be either positive (favoring response A) or negative (favoring response B); (ii) boundary separation, the distance by which the two decision boundaries are separated; (iii) starting point, which captures an initial bias towards one or the other choice (starting point = 0.5 indicates an unbiased decision maker), and (iv) a non-decisional constant ($t_0$), which accounts for net delay due to sensory encoding (before decisional process) and motor execution (after a decision has been made); not illustrated here. **(B)** Exclusion of trials with outlier RT values (inordinately short or long RTs) prior to diffusion modeling (Methods): Response accuracy plotted as a function of RT (binned into 50 ms bins). Here, accuracy was computed by first pooling trials across all mice, all sessions (of 50-50 conditions), and both locations, and then computing % correct from the trials within each RT bin. Gray shading: 95% C.I. obtained by bootstrapping. Based on this data, trials with RTs shorter than 300 ms or longer than 2550 ms, which exhibited accuracy that was not distinguishable from chance, were excluded from diffusion modeling. The same window was then applied to data of 90u-10 condition. **(C)** Outlier trial exclusion did not impact the effects on median RT reported in Fig. 2A (median RT was shorter in the 90u-10 condition at both locations); conventions as in Fig. 2A. **(D-F)** Results from drift diffusion modeling: effect of spatial probability manipulation on the other two parameters of the model - boundary separation (D), and non-decisional constant ($t_0$; E). No systematic modulation of either parameter, at either location (green: upper location; purple: lower location), by spatial probability. Effect of spatial probability on 'threshold to correct responses' (F; a quantity computed as the difference between upper boundary and starting point). Conventions as in C.

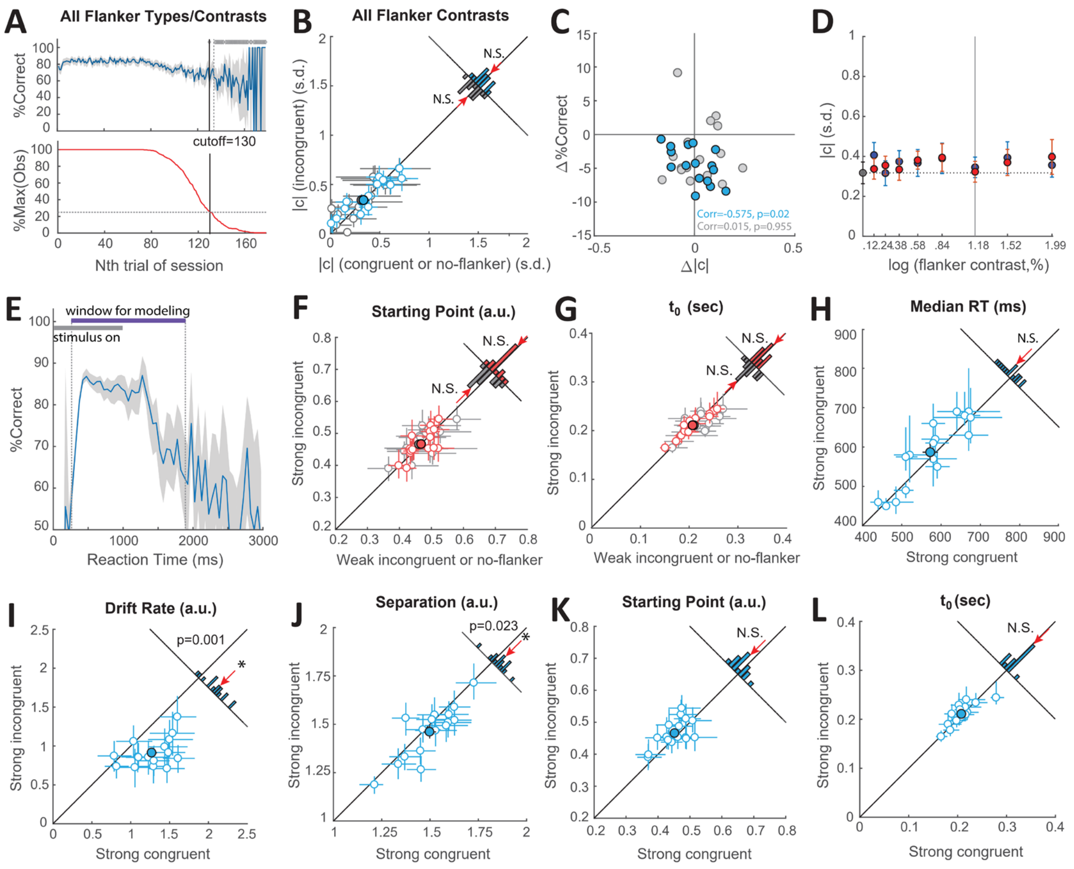


**Figure S3. Related to Figures 3 and 4. Flanker task: trial exclusion criterion, and effect of flanker contrast on RTs and parameters of the drift diffusion model in freely behaving mice.**
**(A)** Identification of trials (late in sessions) that corresponded to animals being poorly engaged in the task (Methods; conventions as in Fig. S1D). *Top panel*: Time course of overall response accuracy across mice (n=16) as a function of trial number within session. Accuracy obtained from trials pooled across all mice, conditions and contrasts, and computed as a function of trial number within session (blue; Methods). *Bottom panel*: Number of actual observations across mice for each trial number, as a percentage of the maximal number of possible observations (mice * sessions), plotted as a function of trial number within session (red). Based on these data, all trials above 130 of each behavioral session of this task were dropped from analysis (Methods). Results in Figures 3 and 4 are based on data from trials 1-130 from each behavioral session. **(B)** Scatter plot comparing response criterion between the flanker conditions; conventions as in Fig. 3E. **(C)** Plots of changes in response accuracy against changes in criterion. Blue: incongruent trials versus congruent trials, gray data: incongruent trials versus no-flanker trials. Corr=Pearson's correlation; **(D)** Plot of decision criterion as a function of flanker contrast. Gray vertical line: fixed contrast of the target. Gray data: No flanker; blue: congruent flanker; red: incongruent flanker. **(E)** Exclusion of trials with outlier RT values (inordinately short or long RTs) prior to diffusion modeling (Methods); all conventions same as in Fig. S2B. Response accuracy plotted as a function of RT (50 ms bins). Based on this data, trials with RTs shorter than 250 ms or longer than 1900 ms, which exhibited accuracy that was not distinguishable from chance, were excluded from diffusion modeling. **(F, G)** Drift diffusion modeling. Scatter plots comparing starting point (F), and non-decisional constant ($t_0$, G) in trials with strong versus weak incongruent flankers (red data), and in trials with strong incongruent flankers versus no flanker (gray data). *Insets:* Distributions of changes in starting point (F) and $t_0$ (G). **(H)** Scatter plot comparing median RTs in trials with strong incongruent versus strong congruent flankers. **(I-L)** Drift diffusion modeling. Scatter plots comparing drift rate (I), boundary separation (J), starting point (K) and non-decisional constant (L), in trials with strong incongruent versus strong congruent flankers. *Insets:* Distributions of changes in corresponding parameters. Incongruent flankers produced a reduction in drift rate (I) as well as a reduction in boundary separation (J) as

compared to the congruent trials, together explaining a lack of increase in overall RT. *: $p<0.05$, N.S.: not significant; signed-rank and HB tests.
